# Airborne Sound Detection in *Manduca sexta* Caterpillars

**DOI:** 10.64898/2026.09.19.752919

**Authors:** Sara Aghazadeh, Aishwarya Sriram, Carol I. Miles, Ronald N. Miles

**Affiliations:** Department of Mechanical Engineering, Binghamton University, Binghamton, NY, USA; Department of Biological Sciences, Binghamton University, Binghamton, NY, USA

**Keywords:** *Manduca sexta*, insect hearing, airborne sound, substrate vibration, mechanosensory hairs, behavioural response

## Abstract

We investigated the hearing mechanism of *Manduca sexta* caterpillars to determine whether their behavioural responses to acoustic stimulation are caused by sound-induced substrate vibration or by direct detection of airborne sound.

Behavioural response thresholds to direct substrate vibration and airborne sound were measured using pure-tone stimuli at 150 Hz and 2000 Hz. A frequency of 150 Hz was selected based on previous behavioural measurements showing the lowest sound detection thresholds between approximately 100 and 200 Hz, while 2000 Hz was selected based on laser Doppler vibrometry measurements that revealed a mechanical resonance near this frequency in a thoracic hair.

At both test frequencies, the direct substrate vibration detection thresholds were more than 100 times greater, on average, than the sound-induced substrate vibrations measured at the behavioural sound detection thresholds. Thus, the behavioural responses to airborne sound cannot be explained by sound-induced substrate vibration.

During airborne sound stimulation, caterpillars exhibited graded behavioural responses with increasing sound pressure level, progressing from freezing to twitching and finally to jump-startle. Together, these findings provide direct experimental evidence that the behavioural responses of *M. sexta* caterpillars to acoustic stimulation cannot be explained by sound-induced substrate vibration and instead support direct detection of airborne sound. Understanding how caterpillars respond to airborne sound may also provide biological inspiration for the development of bio-inspired acoustic sensors.

## 1 Introduction

Understanding how insects detect sound and vibration provides insight into biological mechanosensory systems Yack (2004), Windmill & Jackson (2016) and may also guide the development of bio-inspired acoustic sensors Robert et al. (2010), Miles et al. (2009), Zhang et al. (2018), Zhou & Miles (2017). Many insects use mechanical stimuli for communication and for detecting predators, prey, and other animals in their environment Yack (2016), Casas & Dangles (2010), Raboin & Elias (2019).

Insects and other arthropods have evolved a variety of sensory structures for detecting mechanical stimuli, including tympanal organs, chordotonal organs such as Johnston’s organ, and mechanosensory hairs Yack (2004), Windmill & Jackson (2016). Depending on the species and sensory structure, these systems can detect airborne sound, including sound pressure and air-particle motion, or vibrations transmitted through a substrate Yack (2004), Raboin & Elias (2019), Hill & Wessel (2016). These different forms of mechanical sensing are adapted to the physical characteristics of the signals that are important to each species.

For example, some moths, crickets, and katydids detect airborne sound using tympanal organs, in which sound causes vibration of a thin tympanal membrane Yack (2004), Windmill & Jackson (2016), Hoy & Robert (1996), Montealegre-Z & Robert (2015). Other insects use chordotonal organs to detect mechanical stimuli.

In mosquitoes and some flies, Johnston’s organ in the antenna detects vibrations of the antennal structures associated with airborne sound Göpfert & Robert (2002), Menda et al. (2019). Substrate-borne vibrations can also be detected through specialized sensory organs in the legs.

For example, the southern green stink bug, *Nezara viridula*, uses substrate-borne vibrations for courtship communication and detects these vibrations through sensory organs in its legs Janža et al. (2024).

Termites also use specialized mechanosensory organs in the legs and antennae to detect substrate vibrations Sansom et al. (2025).

In spiders, substrate-borne vibrations can also play an important role in communication; male peacock spiders, *Maratus volans*, produce vibrational signals together with visual displays during courtship Girard et al. (2011).

Mechanosensory hairs provide another means of detecting air motion associated with airborne sound. Spider silk is highly responsive to airborne sound and can closely follow the motion of the surrounding air Zhou & Miles (2017). Orb-weaving spiders can use sound-induced motion of their webs to detect airborne acoustic stimuli Zhou et al. (2022).

Caterpillars provide another interesting example of mechanical sensing. Although they lack the tympanal ears found in many adult insects, several caterpillar species show clear behavioural responses to acoustic stimuli Minnich (1925, 1936), Markl & Tautz (1975). Early studies reported sound-evoked responses in caterpillars, including rapid body movements and startle-like behaviours Minnich (1925, 1936). Similar responses reported across caterpillar species include freezing, twitching, squirming, flicking, and startle displays, many of which are considered defensive responses that may help reduce the risk of predation Myers & Smith (1978), Tautz & Markl (1978), Drinkwater et al. (2022). Later studies of the cabbage moth caterpillar, *Mamestra brassicae*, showed that mechanosensory hairs respond to airborne acoustic stimuli Markl & Tautz (1975), Tautz (1977, 1978, 1979). These hairs were also proposed to help caterpillars detect approaching flying wasps and initiate defensive responses Tautz & Markl (1978).

More recently, Taylor & Yack (2019) showed that monarch butterfly caterpillars, *Danaus plexippus*, respond to pure tones between 50 and 900 Hz and are most sensitive between 100 and 200 Hz, with the lowest mean behavioural threshold at 150 Hz. The caterpillars exhibited several defensive responses, including freezing, contraction, and flicking. Their study also considered whether these responses could result from vibrations induced in the supporting leaf and found no detectable leaf vibration during acoustic stimulation, supporting an airborne rather than substrate-borne origin of the behavioural responses.

The tobacco hornworm, *Manduca sexta*, exhibits similar sound-evoked defensive behaviours. Previous work conducted in our laboratory showed that fourth-instar *M. sexta* caterpillars respond to pure tones between 50 and 1000 Hz, with the lowest behavioural thresholds between 100 and 200 Hz. Iacovazzi (2023). These findings motivated the selection of 150 Hz as one of the test frequencies in the present study.

Although these studies clearly demonstrate that *M. sexta* caterpillars respond to acoustic stimulation, it remains unclear whether these responses result from direct detection of airborne sound or from sound-induced vibration of the supporting substrate. This distinction is important because caterpillars normally remain in contact with plant leaves and stems, and airborne sound can cause these surfaces to vibrate Caldwell (2014). Therefore, two types of substrate vibration must be considered: direct substrate vibration, in which the substrate itself is mechanically driven, and sound-induced substrate vibration, in which airborne sound causes the supporting substrate to vibrate. Comparing direct substrate vibration detection thresholds with sound-induced substrate vibrations measured during acoustic stimulation can determine whether substrate vibration is sufficient to account for the observed behavioural responses.

In the present study, we investigated whether the behavioural responses of fourth-instar *Manduca sexta* caterpillars are elicited by airborne sound or by sound-induced substrate vibration. Behavioural thresholds were measured independently for direct substrate vibration and airborne sound using pure-tone stimuli at 150 Hz and 2000 Hz. The 2000 Hz frequency was selected based on laser Doppler vibrometry measurements that revealed a mechanical resonance near this frequency in a thoracic hair (Supplementary Fig. 6). During airborne sound stimulation, the sound-induced vibration of the supporting substrate was measured and compared with the direct substrate vibration detection thresholds to determine whether substrate vibration was sufficient to account for the observed behavioural responses. We also examined how behavioural responses changed with increasing sound pressure level by classifying the responses as freeze, twitch, or jump-startle. Together, these measurements allowed us to determine whether the observed behavioural responses could be explained by sound-induced substrate vibration or were instead associated with direct detection of airborne sound.

## 2 Materials and Methods

### 2.1 Caterpillar Rearing

*Manduca sexta* caterpillars were obtained from a laboratory colony at Binghamton University and reared individually in containers on a wheat-germ-based artificial diet, following established rearing procedures Yamamoto (1969), del Campo & Miles (2003). Fourth-instar caterpillars were used in all behavioural experiments.

### 2.2 Overview of the Experimental Design

To determine whether the behavioural responses of *Manduca sexta* caterpillars to acoustic stimulation could be explained by sound-induced substrate vibration, two complementary behavioural experiments were performed at 150 and 2000 Hz.

First, direct substrate vibration experiments were performed to determine the behavioural substrate vibration detection threshold. A range of vibration amplitudes was applied while the substrate acceleration was measured and the caterpillars’ behavioural responses were recorded. Because the geometry and mechanical properties of natural leaves and stems can vary substantially, an electrodynamic shaker was used as a controlled and repeatable substrate for these experiments. Second, airborne sound experiments were performed using pure-tone stimuli over a range of sound pressure levels. During these experiments, the sound pressure level and the sound-induced vibration of the supporting substrate were measured while the caterpillars’ behavioural responses were recorded.

The direct substrate vibration detection thresholds were then compared with the sound-induced substrate vibrations measured at the behavioural sound detection thresholds to determine whether the substrate vibrations generated during airborne sound stimulation were sufficient to elicit the observed behavioural responses.

### 2.3 Direct Substrate Vibration Experimental Setup

The direct substrate vibration experiment was conducted to determine the behavioural detection threshold of fourth-instar *Manduca sexta* caterpillars to direct substrate vibration.

Direct substrate vibrations were generated using a B&K Type 4809 electrodynamic vibration exciter (Brüel & Kjær, Denmark), as shown in Fig. 1(a). Sinusoidal excitation signals were generated in MATLAB and output through a National Instruments (NI) PXI-1033 data acquisition system. The output signal was amplified using a Crown D-75 power amplifier before driving the vibration exciter.

**Figure 1:**
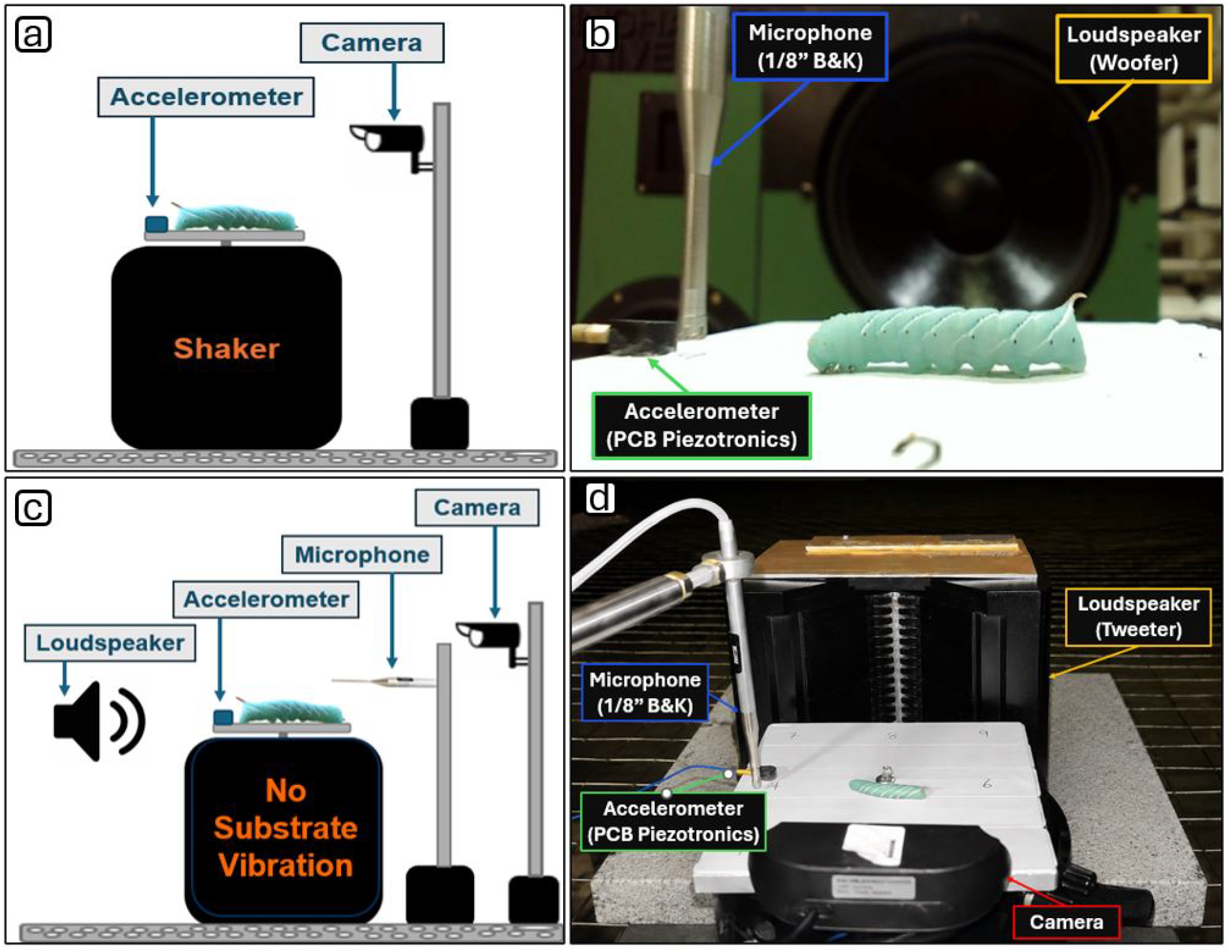
Experimental setups for direct substrate vibration and airborne sound stimulation. (a) Schematic of the direct substrate vibration experiment. Substrate vibrations were generated using a Brüel & Kjær (B&K) Type 4809 electrodynamic shaker, a PCB Piezotronics Model 352A24 accelerometer measured the applied vibration, and a digital camera recorded the caterpillar’s behavioural responses. (b) Schematic of the airborne sound experiment. The shaker was not mechanically driven and served only as the supporting substrate. Pure-tone stimuli were generated by a loudspeaker, a B&K Type 4138 1/8-inch microphone measured the sound pressure level, and the accelerometer measured sound-induced substrate vibration. (c) Experimental arrangement for airborne sound stimulation at 150 Hz in the anechoic chamber using a woofer. (d) Experimental arrangement for airborne sound stimulation at 2000 Hz using a tweeter.

A PCB Piezotronics Model 352A24 shear accelerometer (PCB Piezotronics Inc., USA; sensitivity: 92.8 mV/g) was mounted adjacent to the caterpillar on the shaker platform using a thin layer of mounting wax. The accelerometer continuously measured the applied substrate vibration, and its output was acquired using the NI PXI-1033 data acquisition system. A digital camera positioned near the shaker recorded the caterpillar’s behavioural responses throughout each experiment.

Twenty fourth-instar *Manduca sexta* caterpillars (N=20) were tested, with ten individuals (n=10) assigned to each frequency. Each caterpillar was placed individually on the shaker platform and exposed to sinusoidal substrate vibrations spanning acceleration amplitudes below and above the expected behavioural threshold. To minimize possible effects of habituation or fatigue, five caterpillars at each frequency received the stimuli in ascending order of amplitude, whereas the remaining five received the same stimuli in descending order. The applied substrate vibration, expressed as acceleration (m/s^2^), was recorded by the accelerometer for every stimulus level.

The behavioural detection threshold was defined as the lowest stimulus amplitude that elicited an observable behavioural response in at least two of three repeated trials at that stimulus amplitude.

Behavioural responses were classified by visual inspection of the video recordings for each stimulus presentation as: (1) freeze, characterized by cessation of body movement; (2) twitch, consisting of a localized movement of one or more body segments; and (3) jump-startle, characterized by rapid lifting of the thoracic and anterior abdominal segments. Freeze represented the lowest-level behavioural response.

A continuous whole-body tremor was also observed during some direct substrate vibration trials, particularly at amplitudes between those associated with twitch and jump-startle responses. During these trials, the continuous body oscillation made it difficult to determine whether twitch or jump-startle responses also occurred. Tremor was therefore documented separately in Fig. 2(a,b) but was not classified as a behavioural response because it may have resulted from direct mechanical forcing by the shaker. Tremor was not considered when determining the behavioural detection threshold.

**Figure 2:**
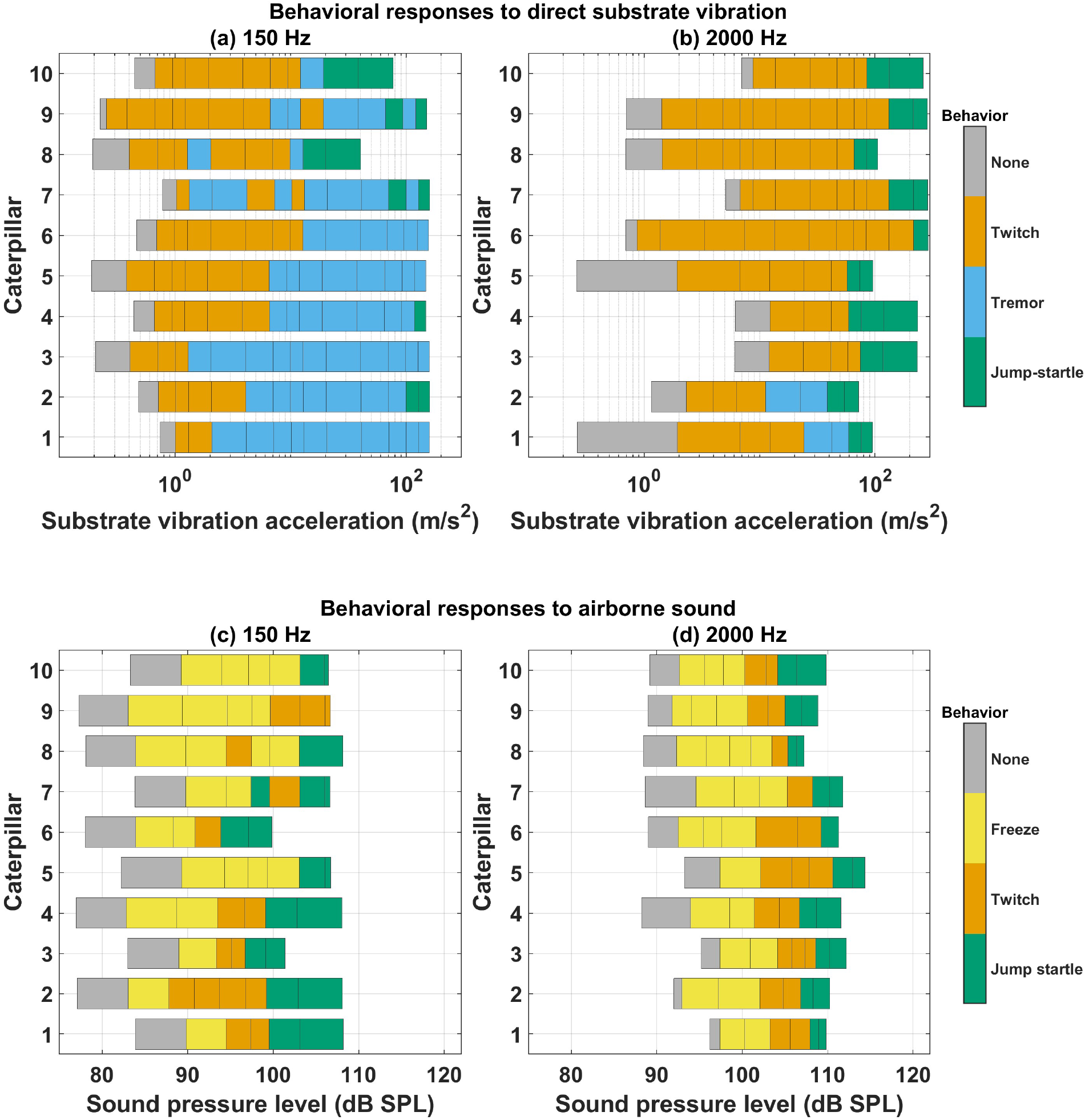
Behavioural responses of fourth-instar *Manduca sexta* caterpillars to direct substrate vibration and airborne sound. (a,b) Responses measured during direct substrate vibration at 150 Hz and 2000 Hz, respectively. (c,d) Responses measured during airborne sound stimulation at 150 Hz and 2000 Hz, respectively. Each horizontal row represents one caterpillar, and the colored segments indicate the observed response over the corresponding range of substrate vibration acceleration in (a,b) or sound pressure level in (c,d). Tremor denotes continuous whole-body oscillation observed during direct substrate vibration and is shown to document the observed motion; it was not classified as a behavioural response because it may have resulted from direct mechanical forcing by the shaker. Independent groups of ten caterpillars were tested at each frequency.

### 2.4 Airborne Sound Experimental Setup

The airborne sound experiment was conducted to determine the behavioural sound detection threshold of fourth-instar *Manduca sexta* caterpillars while simultaneously measuring the sound-induced vibration of the supporting substrate generated by the incident sound field.

The same experimental platform used for the direct substrate vibration experiments was employed to maintain identical contact conditions between the caterpillar and the substrate. During the airborne sound experiments, however, the B&K Type 4809 electrodynamic shaker remained stationary and served only as the supporting substrate; no mechanical excitation was applied.

Figure 1(b) illustrates the instrumentation and measurement configuration used for the airborne sound experiments. Pure-tone stimuli at 150 Hz and 2000 Hz were generated in MATLAB and output through a National Instruments (NI) PXI-1033 data acquisition system. The signals were amplified using Crown-DC-75 amplifier before being delivered to the loudspeakers. A woofer was used for the 150 Hz experiments, whereas a tweeter was used for the 2000 Hz experiments.

The loudspeaker-to-caterpillar distance was selected to maintain the caterpillar within the acoustic near field while providing a sufficient sound level. For the 150 Hz experiments, the woofer was positioned approximately 50 cm from the caterpillar (acoustic wavelength *≈* 2.3 m), whereas for the 2000 Hz experiments, the tweeter was positioned approximately 14 cm from the caterpillar (acoustic wavelength *≈* 17 cm).

A calibrated B&K Type 4138 (1/8-inch) reference microphone was positioned adjacent to the caterpillar to measure the sound pressure level (SPL) at the caterpillar’s location. A PCB Piezotronics Model 352A24 shear accelerometer was mounted on the stationary shaker platform adjacent to the caterpillar using a thin layer of mounting wax to measure the sound-induced substrate vibration. The microphone and accelerometer signals were simultaneously acquired using the NI PXI-1033 data acquisition system. The experimental arrangements used for the 150 Hz and 2000 Hz airborne sound experiments are shown in Fig. 1(c,d).

Twenty fourth-instar *M. sexta* caterpillars (*N* = 20) were tested, with ten individuals (*n* = 10) assigned to each frequency. Each caterpillar was exposed to pure-tone stimuli over a range of sound pressure levels spanning values below and above the expected behavioural detection threshold. For each stimulus presentation, the sound pressure level, corresponding sound-induced substrate vibration expressed as acceleration (m/s^2^), and behavioural response were recorded simultaneously. The behavioural sound detection threshold was defined as the lowest sound pressure level that elicited an observable behavioural response in at least two of three repeated trials at that sound pressure level.

Behavioural responses were classified by visual inspection of the video recordings for each stimulus presentation as: (1) none, defined as no observable response or change in behaviour; (2) freeze, characterized by cessation of ongoing movement; (3) twitch, consisting of localized movement of one or more body segments; and (4) jump-startle, characterized by rapid lifting of the thoracic and anterior abdominal segments. Representative examples of freeze, twitch, and jump-startle responses are provided in Supplementary Movies S1-S6.

For each caterpillar, the sound-induced substrate vibration measured at the behavioural sound detection threshold was subsequently compared with the direct substrate vibration detection threshold, with both quantities expressed as acceleration (m/s^2^). This comparison was used to determine whether sound-induced substrate vibration was sufficient to account for the behavioural responses observed during airborne sound stimulation.

### 2.5 Statistical Analysis

Nonparametric statistical comparisons were performed using the Mann-Whitney *U* test. Sound pressure level (SPL) thresholds for freeze, twitch, and jump-startle responses were compared between 150 Hz and 2000 Hz. Sound-induced substrate vibration amplitudes measured at the behavioural sound detection thresholds were also compared between the two frequencies. In addition, at each frequency, the direct substrate vibration detection thresholds were compared with the sound-induced substrate vibration amplitudes measured at the behavioural sound detection thresholds. Statistical significance was defined as *P <* 0.05. Statistical analyses were performed using MATLAB (MathWorks, Natick, MA, USA) and Microsoft Excel.

## 3 Results

### 3.1 Direct Substrate Vibration Experiment Results

#### 3.1.1 Behavioural Responses to Substrate Vibration

The behavioural responses of fourth-instar *Manduca sexta* caterpillars to direct substrate vibration were examined at 150 Hz and 2000 Hz. Figure 2(a,b) shows the behavioural responses at the two test frequencies. Each horizontal row represents an individual caterpillar, and the colored segments indicate the observed response over the corresponding range of substrate vibration accelerations.

#### 3.1.2 Direct Substrate Vibration Detection Threshold

At 150 Hz, the direct substrate vibration detection threshold ranged from 0.27 to 1.19 m/s^2^, with a median of 0.825 m/s^2^ and a mean (*±* SD) of 0.772 *±* 0.288 m/s^2^. At 2000 Hz, the threshold ranged from 0.98 to 17.31 m/s^2^, with a median of 5.259 m/s^2^ and a mean (*±* SD) of 7.088*±*5.972 m/s^2^. The direct substrate vibration detection threshold was significantly higher at 2000 Hz than at 150 Hz (Mann-Whitney *U* test, *P* = 0.00033), indicating lower sensitivity to direct substrate vibration at 2000 Hz than at 150 Hz.

These direct substrate vibration detection thresholds were used as reference values for comparison with the sound-induced substrate vibrations measured during the airborne sound experiments, as shown later in Fig. 4.

### 3.2 Airborne Sound Experiment Results

#### 3.2.1 Behavioural Responses to Airborne Sound

The behavioural responses of fourth-instar *Manduca sexta* caterpillars to airborne sound were examined at 150 Hz and 2000 Hz. Figure 2(c,d) shows the behavioural responses at the two test frequencies. Each horizontal row represents an individual caterpillar, and the colored segments indicate the observed behavioural response over the corresponding range of sound pressure levels (SPLs).

At both frequencies, caterpillars exhibited a graded progression of behavioural responses as the sound pressure level increased. No observable behavioural response occurred at the lowest sound pressure levels. As the sound pressure level increased, caterpillars first exhibited freezing, followed by twitching and finally jump-startle. Although the sequence of responses was similar at both frequencies, higher sound pressure levels were generally required to elicit each response at 2000 Hz than at 150 Hz.

Representative examples of freezing, twitching, and jump-startle behaviours observed during airborne sound stimulation are provided in Supplementary Movies S1-S6.

The SPL required to elicit each behavioural response was significantly higher at 2000 Hz than at 150 Hz (Fig. 3; Table 1).

**Table 1:** Airborne sound pressure level thresholds for individual behavioural responses at 150 Hz and 2000 Hz. Values are reported as mean *±* SD. Frequencies were compared using the Mann-Whitney *U* test. Sample sizes were *n* = 10 at both frequencies for freeze, *n* = 8 at 150 Hz and *n* = 10 at 2000 Hz for twitch, and *n* = 9 at 150 Hz and *n* = 10 at 2000 Hz for jump-startle.

| Behaviour | 150 Hz (dB SPL) | 2000 Hz (dB SPL) | $P$-value |
| --- | --- | --- | --- |
| Freeze | $89 \pm 3$ | $96 \pm 2$ | 0.0002 |
| Twitch | $95 \pm 3$ | $103 \pm 1$ | $< 0.0001$ |
| Jump-startle | $101 \pm 3$ | $108 \pm 2$ | 0.0002 |

**Figure 3:**
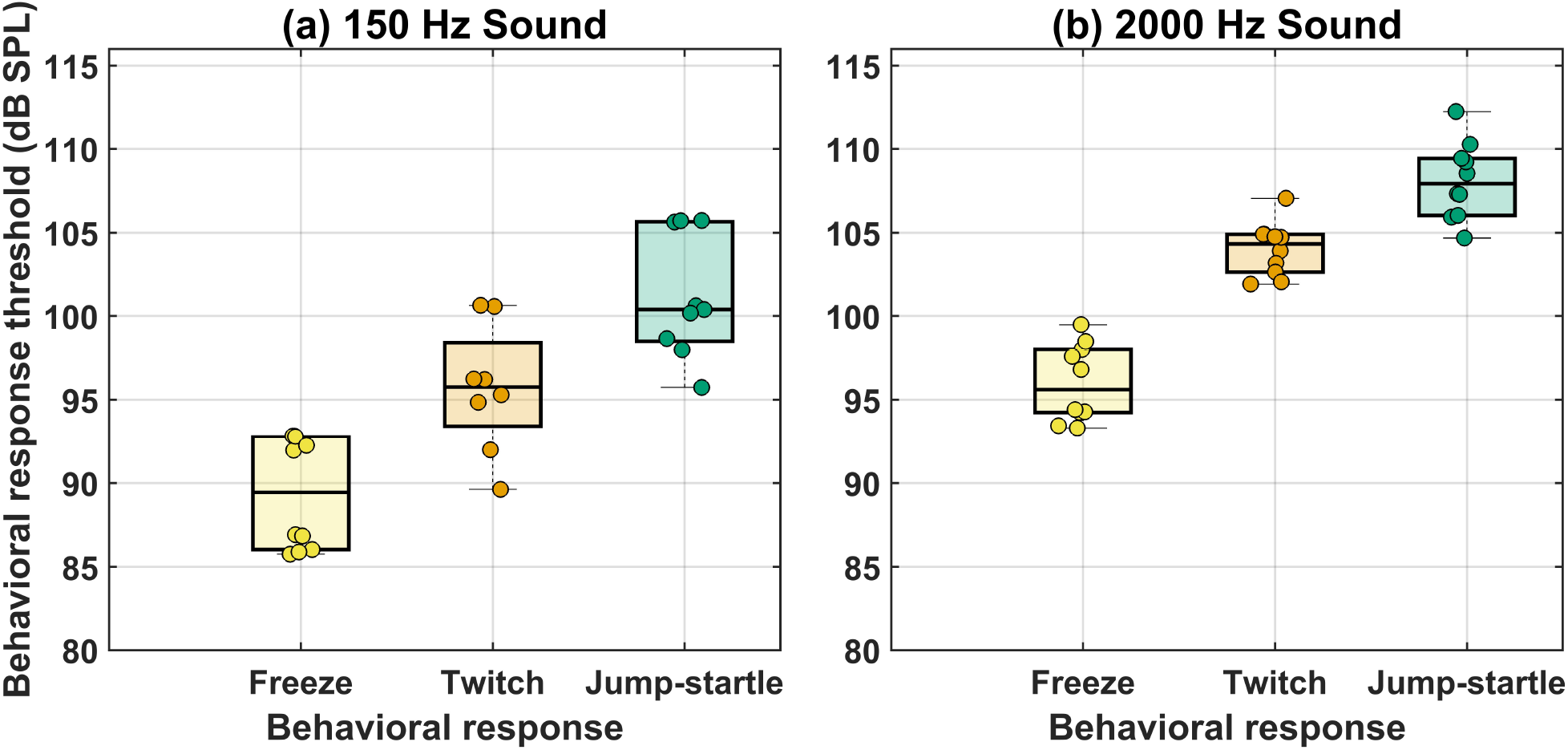
Different sound pressure levels elicited different behavioural responses at 150 Hz and 2000 Hz. Thresholds are shown for freeze, twitch, and jump-startle responses. Individual points represent individual caterpillars, boxes indicate the interquartile range, horizontal lines indicate the median, and whiskers indicate the data range.

**Figure 4:**
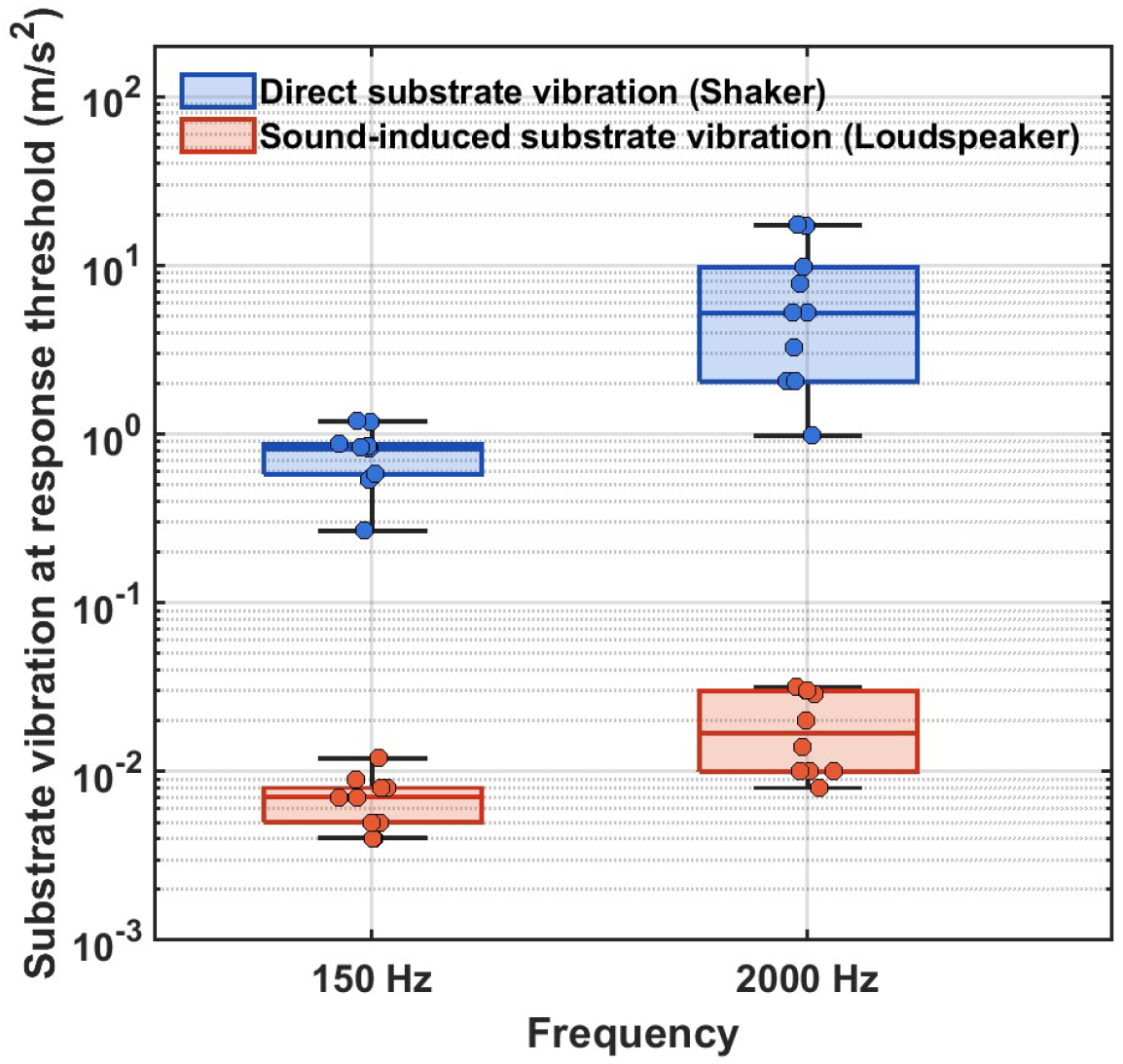
Comparison of the direct substrate vibration detection thresholds and the sound-induced substrate vibrations measured at the behavioural sound detection thresholds. Each point represents one caterpillar. Boxes indicate the median and interquartile range (25th-75th percentiles), and whiskers indicate the minimum and maximum values. At both frequencies, the direct substrate vibration detection thresholds were significantly greater than the corresponding sound-induced substrate vibrations (Mann-Whitney *U* test, *P <* 0.0001).

Freeze thresholds increased from 89*±*3 dB SPL (mean *±* SD; *n* = 10) at 150 Hz to 96*±*2 dB SPL (*n* = 10) at 2000 Hz (Mann-Whitney *U* test, *P* = 0.0002). Similarly, twitch thresholds increased from 95 *±* 3 dB SPL (*n* = 8) to 103 *±* 1 dB SPL (*n* = 10) (*P <* 0.0001), and jump-startle thresholds increased from 101 *±* 3 dB SPL (*n* = 9) to 108 *±* 2 dB SPL (*n* = 10) (*P* = 0.0002). The smaller sample sizes for twitch and jump-startle at 150 Hz reflect individuals that did not exhibit these responses within the tested sound pressure level range. At both frequencies, progressively stronger behavioural responses required progressively higher sound pressure levels, demonstrating a graded behavioural response to airborne sound.

#### 3.2.2 Behavioural Sound Detection Threshold

At 150 Hz, the behavioural sound detection threshold ranged from 86 to 93 dB SPL, with a median of 89 dB SPL and a mean (*±* SD) of 89 *±* 3 dB SPL. At 2000 Hz, the threshold ranged from 93 to 99 dB SPL, with a median of 96 dB SPL and a mean (*±* SD) of 96 *±* 2 dB SPL. The behavioural sound detection threshold was significantly higher at 2000 Hz than at 150 Hz (Mann-Whitney *U* test, *P* = 0.0002).

These behavioural sound detection thresholds were used to identify the corresponding sound-induced substrate vibrations measured during airborne sound stimulation.

#### 3.2.3 Sound-Induced Substrate Vibration at the Behavioural Sound Detection Threshold

For each caterpillar, the sound-induced substrate vibration corresponding to the behavioural sound detection threshold was determined from the acceleration measured during airborne sound stimulation.

At 150 Hz, the sound-induced substrate vibration ranged from 0.004 to 0.012 m/s^2^, with a median of 0.0075 m/s^2^ and a mean (*±*SD) of 0.0069 *±*0.0025 m/s^2^. At 2000 Hz, the sound-induced substrate vibration ranged from 0.008 to 0.032 m/s^2^, with a median of 0.0170 m/s^2^ and a mean (*±* SD) of 0.0193 *±* 0.0100 m/s^2^. The sound-induced substrate vibration at the behavioural sound detection threshold was significantly greater at 2000 Hz than at 150 Hz (Mann-Whitney *U* test, *P* = 0.0038).

These sound-induced substrate vibration amplitudes were subsequently compared with the direct substrate vibration detection thresholds, as shown in Fig. 4.

### 3.3 Comparison of Direct and Sound-Induced Substrate Vibrations

Figure 4 compares the direct substrate vibration detection thresholds with the sound-induced substrate vibrations measured at the behavioural sound detection thresholds during the airborne sound experiments.

At both frequencies, the sound-induced substrate vibrations were substantially lower than the direct substrate vibration detection thresholds, with no overlap between the distributions (Fig. 4). At 150 Hz, the mean direct substrate vibration detection threshold was approximately 112 times greater than the mean sound-induced substrate vibration measured at the behavioural sound detection threshold (0.772 versus 0.0069 m/s^2^). At 2000 Hz, the mean direct substrate vibration detection threshold was approximately 367 times greater than the mean sound-induced substrate vibration (7.088 versus 0.0193 m/s^2^). Within each frequency, the direct substrate vibration detection thresholds were significantly greater than the sound-induced substrate vibrations (Mann-Whitney *U* test, *P <* 0.0001).

These results demonstrate that the substrate vibrations generated by airborne sound were far below the vibration amplitudes required to elicit behavioural responses during direct substrate vibration. Therefore, the behavioural responses observed during airborne sound stimulation cannot be explained by sound-induced substrate vibration and instead support direct detection of airborne sound by fourth-instar *Manduca sexta* caterpillars.

## 4 Discussion

The primary objective of this study was to determine whether the behavioural responses of fourth-instar *Manduca sexta* caterpillars are elicited by airborne sound or by sound-induced substrate vibration. Our results show that the sound-induced substrate vibrations measured at the behavioural sound detection thresholds were far below the direct substrate vibration amplitudes required to elicit behavioural responses. The complete absence of overlap between the direct substrate vibration detection thresholds and the sound-induced substrate vibrations at both frequencies provides strong evidence that sound-induced substrate vibration cannot account for the observed behavioural responses. Therefore, the behavioural responses observed during airborne sound stimulation are most consistent with direct detection of airborne sound.

Our findings extend previous behavioural observations showing that *Manduca sexta* caterpillars respond to low-frequency airborne sound, with their greatest sensitivity near 150 Hz Iacovazzi (2023). Similar sound-evoked behavioural responses have been reported in other caterpillar species. In monarch butterfly caterpillars, *Danaus plexippus*, Taylor & Yack (2019) found no detectable vibration of the supporting leaf during acoustic stimulation, supporting detection of airborne sound rather than substrate-borne vibration. Earlier studies of *Mamestra brassicae* also demonstrated that mechanosensory hairs respond to airborne acoustic stimuli and may mediate defensive responses to approaching flying wasps Markl & Tautz (1975), Tautz (1977, 1978, 1979), Tautz & Markl (1978). By independently measuring the direct substrate vibration detection thresholds and the sound-induced substrate vibrations during airborne sound stimulation, the present study extends these previous observations by providing quantitative evidence that the sound-induced substrate vibrations were insufficient to account for the observed behavioural responses.

Laser Doppler vibrometry measurements revealed a mechanical resonance near 2000 Hz in a thoracic mechanosensory hair (Supplementary Fig. 6), providing the basis for including 2000 Hz as one of the test frequencies. Although the mean direct substrate vibration detection threshold at 2000 Hz (7.09 m s^*−*2^) appears relatively large when expressed as acceleration, this value corresponds to a displacement amplitude of approximately 45 nm. For sinusoidal motion, acceleration and displacement amplitudes are related by *a* = *ω*^2^*x*, where *ω* = 2*πf*. Thus, at high frequencies, even nanometer-scale substrate displacements can correspond to relatively large acceleration amplitudes. The behavioural experiments also revealed a graded response to airborne sound levels. Lower sound pressure levels predominantly elicited freezing, intermediate levels elicited twitching, and higher levels elicited jump-startle responses. The distinct SPL thresholds for these responses indicate that *Manduca sexta* caterpillars do not exhibit a simple all-or-none response to airborne sound, but instead show progressively stronger behavioural responses as sound level increases.

Similar defensive responses have been reported in other caterpillar species and have been interpreted as antipredator behaviours Myers & Smith (1978), Tautz & Markl (1978), Taylor & Yack (2019), Drinkwater et al. (2022).

Although the present study did not directly investigate predator interactions, the graded behavioural responses observed here may have ecological significance. Mechanosensory hairs in caterpillars have previously been proposed to detect airborne vibrations generated by approaching predatory wasps Tautz & Markl (1978). Whether the progression from freeze to twitch and jump-startle observed in *Manduca sexta* reflects changes in predator proximity or threat level remains to be tested experimentally.

Beyond its biological significance, understanding how caterpillars detect and respond to airborne mechanical stimuli may also have implications for the development of bio-inspired acoustic sensors. The present study demonstrates that the behavioural responses to airborne sound cannot be explained by sound-induced substrate vibration, but it does not determine which component of the airborne acoustic field is detected by the caterpillars or which sensory receptors are involved. Further studies are needed to distinguish the roles of sound pressure and air-particle motion and to identify the sensory structures responsible for airborne sound detection in *Manduca sexta*.

## 5 Conclusion

This study demonstrates that fourth-instar *Manduca sexta* caterpillars respond to airborne sound at both 150 Hz and 2000 Hz. Behavioural responses were graded according to sound levels, progressing from freezing to twitching and finally to jump-startle as sound pressure level increased.

The sound-induced substrate vibrations measured at the behavioural sound detection thresholds were, on average, more than 100 times smaller than the direct substrate vibration amplitudes required to elicit behavioural responses, with no overlap between the two distributions at either frequency. These findings demonstrate that the behavioural responses observed during airborne sound stimulation cannot be explained by sound-induced substrate vibration and instead support direct detection of airborne sound by *M. sexta* caterpillars.

## Acknowledgements

The authors thank Dr. Jian Zhou, Dr. Junpeng Lai, and Dr. Joseph Cosmo Iacovazzi for their valuable discussions and assistance with the experiments.

## Author Contributions

Sara Aghazadeh and Aishwarya Sriram contributed equally to this work and share first authorship. They jointly conceived and designed the study, developed the experimental setups and protocols, performed the laser vibrometry, mechanical vibration, airborne sound, and behavioural experiments, analyzed and interpreted the data, and wrote the manuscript.

Ronald N. Miles supervised the engineering aspects of the project, contributed to the experimental design and interpretation of the engineering results, and critically revised the manuscript. Carol Miles supervised the biological aspects of the project, contributed to the experimental design and biological interpretation of the results, and critically revised the manuscript. All authors discussed the results, contributed to the interpretation of the findings, reviewed the manuscript, and approved the final version.

## Supplementary Material

### Sensory Structure Analysis Using Laser Doppler Vibrometry

Laser Doppler vibrometry was used to characterize the mechanical response of selected sensory structures on fourth-instar *Manduca sexta* caterpillars. Because mechanosensory hairs in insects can respond to air motion Tautz (1977, 1978, 1979), we investigated whether similar structures in *M. sexta* exhibited measurable mechanical responses to airborne sound. These measurements were used to guide the selection of frequencies for the behavioural experiments. Laser Doppler vibrometry has previously been used to characterize the response of lightweight biological structures to airborne acoustic excitation Zhou et al. (2022).

Two complementary measurements were performed: thermal noise measurements and acoustic frequency response measurements. Thermal noise measurements were used to identify the natural resonance frequencies of the selected sensory structures in the absence of external acoustic stimulation, whereas acoustic frequency response measurements were used to characterize their mechanical response to airborne sound Miles (2024), Lai et al. (2024).

Additional details of thermal-noise and acoustic frequency-response measurement procedures for small acoustic sensing structures are described in Lai et al. (2022, 2024).

All measurements were conducted in the anechoic chamber in the Department of Mechanical Engineering at Binghamton University to minimize acoustic reflections and environmental vibrations. Prior to measurement, fourth-instar *M. sexta* caterpillars were euthanized by freezing for approximately 4 h and then allowed to thaw for approximately 10 min until the body regained sufficient flexibility for mounting. Each caterpillar was mounted vertically on a support rod while leaving the selected sensory structures unconstrained.

A Polytec OFV-534 laser Doppler vibrometer (Polytec GmbH, Germany) was focused near the tip of the selected sensory structure using motorized translation stages. The vibration velocity measured by the laser vibrometer was acquired using a National Instruments PXI-1033 data acquisition system and processed in MATLAB.

For the thermal noise measurements, no external acoustic stimulus was applied. Small mechanical fluctuations of the selected sensory structure were recorded by the laser vibrometer and analyzed using power spectral density (PSD) analysis. The resulting PSD spectra were used to identify the natural resonance frequencies of the selected structures Lai et al. (2024).

For the acoustic frequency response measurements, the selected sensory structure was exposed to controlled pure-tone stimuli over frequencies from 500 Hz to 5 kHz. Stimulus signals were generated in MATLAB and output through the National Instruments PXI-1033 data acquisition system. The analog output signal was passed through a dbx Model 234xs electronic crossover to separate the signal into low-, mid-, and high-frequency bands. The signals were amplified using Crown D-75 and Techron 5530 power amplifiers before driving a woofer, mid-range loudspeaker, and tweeter, respectively. The loudspeaker system was positioned approximately 3 m from the caterpillar.

A calibrated Brüel & Kjær Type 4138 (1/8-inch) reference microphone was positioned adjacent to the measured sensory structure to measure the local sound pressure. The microphone signal was conditioned using a Brüel & Kjær Type 5935L dual microphone power supply and acquired using the National Instruments PXI-1033 data acquisition system. Because the measurements were conducted sufficiently far from the loudspeaker in the anechoic chamber, the incident acoustic field at the measurement location was approximated as a plane wave Miles (2024), Lai et al. (2024).

Under the plane-wave approximation, the air-particle velocity was calculated from the measured sound pressure as Miles (2024), Kinsler et al. (2000)

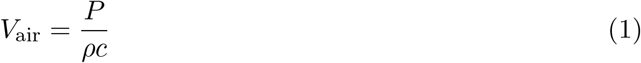

where *P* is the measured sound pressure, *ρ* is the air density, *c* is the speed of sound in air, and *ρc* = 415 kg m^*−*2^ s^*−*1^ is the characteristic acoustic impedance of air.

The mechanical response of each sensory structure to the incident acoustic field was quantified using the normalized velocity ratio Zhou et al. (2022), Lai et al. (2024)

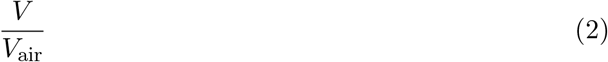

where *V* is the vibration velocity of the sensory structure measured by the laser vibrometer and *V*_air_ is the calculated air-particle velocity. A normalized velocity ratio approaching unity indicates that the vibration velocity of the structure approaches that of the surrounding air-particle motion. This normalization accounts for differences in the applied acoustic stimulus and allows the mechanical response to be compared across frequency.

## Supplementary Results

### Mechanical Response of a Thoracic Mechanosensory Hair

Laser Doppler vibrometry was used to characterize the mechanical response of a representative thoracic mechanosensory hair. Thermal noise measurements were first used to identify its natural resonance, followed by acoustic frequency response measurements to characterize its response to airborne sound.

Figure 6(a) shows the thermal noise PSD measured from a representative thoracic hair. A prominent resonance was observed near 2000 Hz, indicating a natural mechanical resonance of the hair near this frequency.

**Figure 5:**
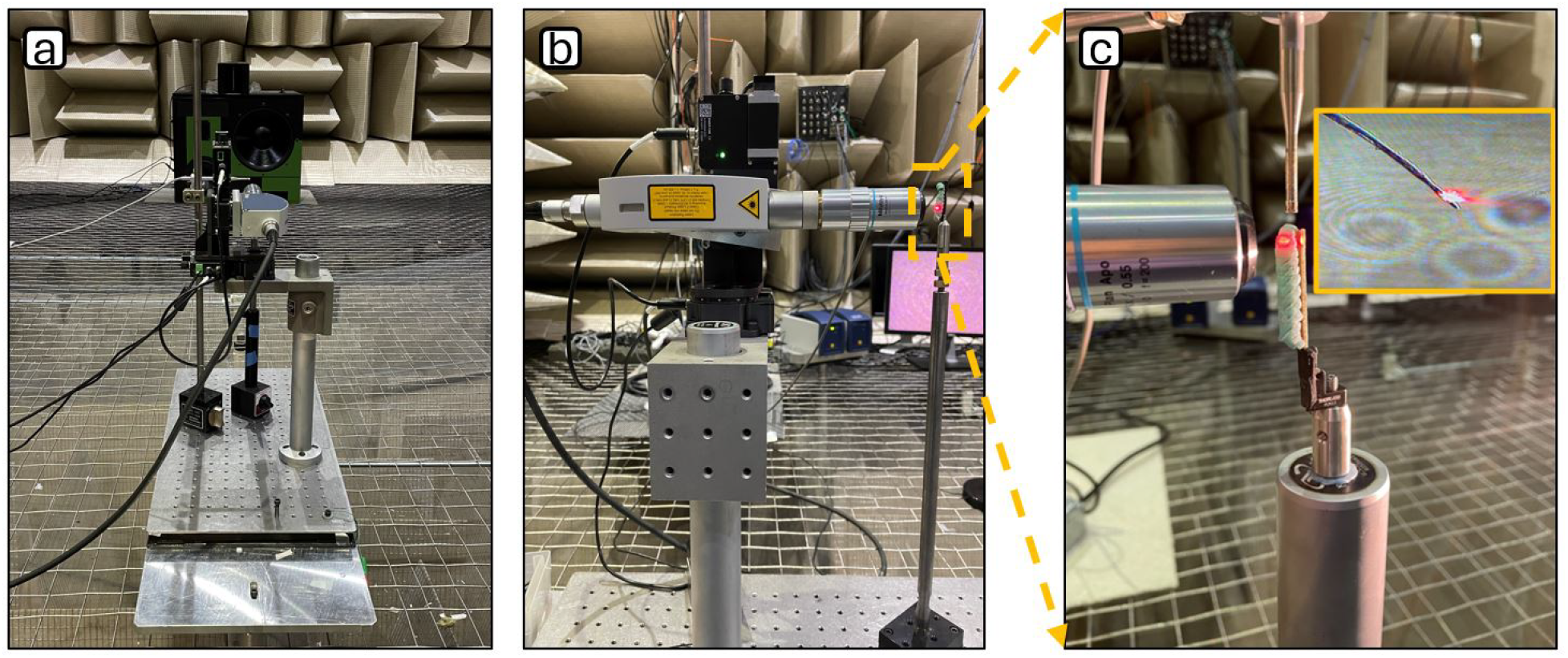
Experimental setup for laser Doppler vibrometry measurements of sensory structures on fourth-instar *Manduca sexta* caterpillars. (a) Side view and (b) front view of the measurement system. The caterpillar was mounted vertically on a support rod, and a Polytec OFV-534 laser Doppler vibrometer measured the vibration velocity of selected sensory structures. (c) Enlarged view of a thoracic hair with the laser beam focused near its tip. The same configuration was used for thermal noise and acoustic frequency response measurements.

**Figure 6:**
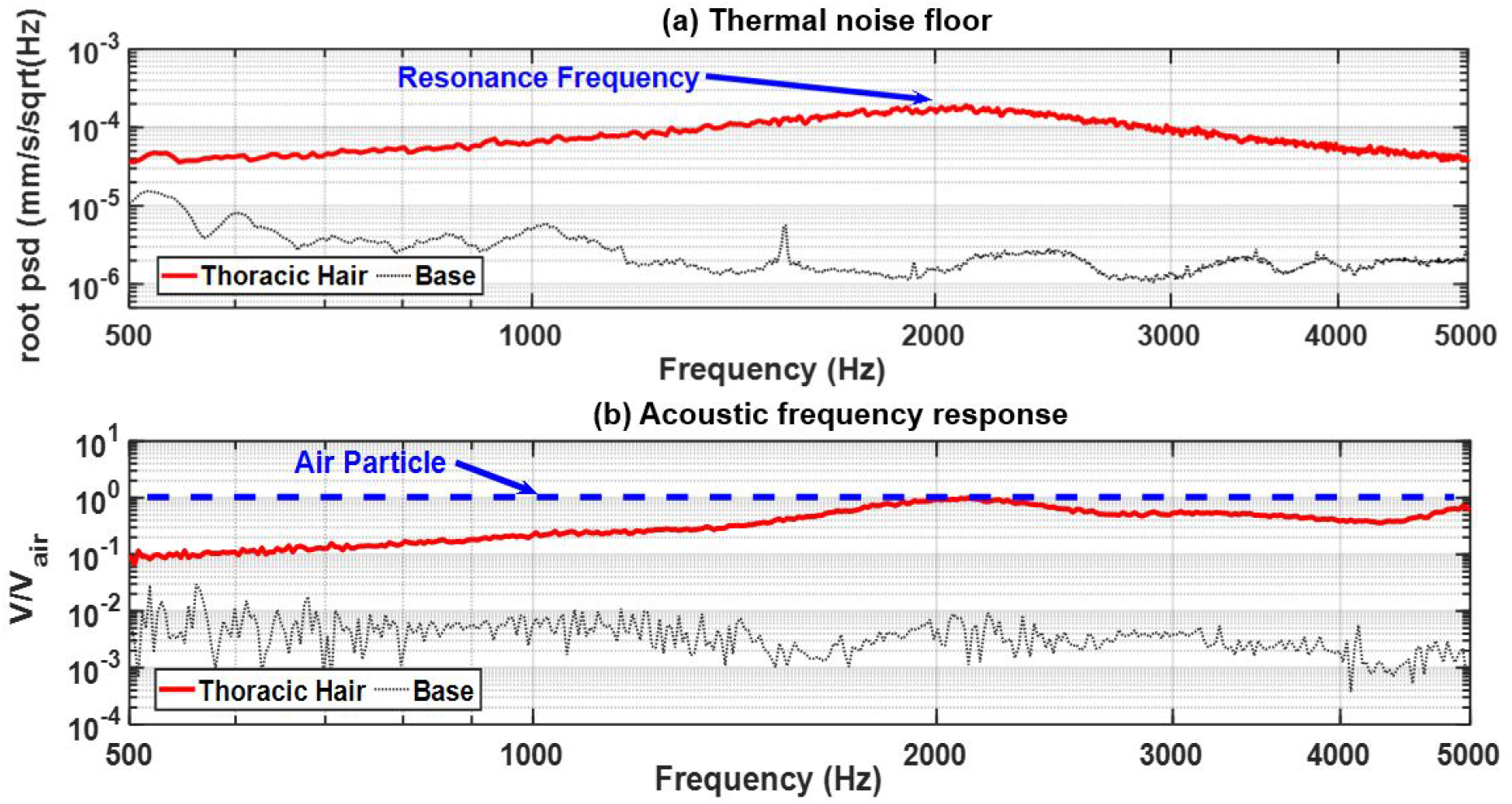
Mechanical characterization of a representative thoracic mechanosensory hair using laser Doppler vibrometry. (a) Power spectral density (PSD) of the hair vibration measured in the absence of external acoustic stimulation. A resonance peak occurs near 2000 Hz. The black curve represents the vibration measured at the supporting base. (b) Acoustic frequency response of the same thoracic hair. The normalized velocity ratio (*V/V*_air_) compares the vibration velocity of the hair measured by laser Doppler vibrometry with the air-particle velocity calculated from the reference microphone measurement. The response approaches unity near 2000 Hz, indicating that the vibration velocity of the hair approaches the surrounding air-particle velocity at this frequency.

The acoustic frequency response of the same thoracic hair is shown in Fig. 6(b). The normalized velocity ratio (*V/V*_air_) increased with frequency and approached unity near 2000 Hz, indicating that the vibration velocity of the hair approached the surrounding air-particle velocity near its mechanical resonance.

Based on these measurements, together with previous behavioural measurements showing strong sensitivity of *M. sexta* caterpillars near 150 Hz Iacovazzi (2023), 150 Hz and 2000 Hz were selected as the two frequencies for the behavioural experiments.

## Supplementary Movies

### Supplementary Movie S1

Representative freeze response of a fourth-instar *Manduca sexta* caterpillar during airborne sound stimulation at 150 Hz.

### Supplementary Movie S2

Representative twitch response of a fourth-instar *Manduca sexta* caterpillar during airborne sound stimulation at 150 Hz.

### Supplementary Movie S3

Representative jump-startle response of a fourth-instar *Manduca sexta* caterpillar during airborne sound stimulation at 150 Hz.

### Supplementary Movie S4

Representative freeze response of a fourth-instar *Manduca sexta* caterpillar during airborne sound stimulation at 2000 Hz.

### Supplementary Movie S5

Representative twitch response of a fourth-instar *Manduca sexta* caterpillar during airborne sound stimulation at 2000 Hz.

### Supplementary Movie S6

Representative jump-startle response of a fourth-instar *Manduca sexta* caterpillar during airborne sound stimulation at 2000 Hz.

